# Stromal Heterogeneity in the Proliferative Endometrial *Functionalis* - A single-cell approach

**DOI:** 10.1101/2020.07.17.203844

**Authors:** Suzanna Queckbörner, Carolina von Grothusen, Nageswara Boggavarapu, Lindsay C. Davies, Kristina Gemzell-Danielsson

## Abstract

The endometrium undergoes regular regeneration and stromal proliferation as part of the normal menstrual cycle. To better understand cellular interactions driving the mechanisms in endometrial regeneration we employed single-cell RNA sequencing. Endometrial samples were obtained during the proliferative phase of the menstrual cycle from healthy women aged 24–32 years. Within the stromal compartment multiple stromal populations were found, suggestive of specific stromal niches that control inflammation and extracellular matrix composition. Ten different stromal cell and two pericyte subsets were identified. Applying different R packages (Seurat, SingleR, Velocyto) we determined cell cluster diversity and cell lineage/trajectory while using external data to validate our findings. By understanding healthy regeneration in the described stromal compartments, we aim to identify points of intervention for novel therapy development in order to treat benign gynaecological disorders affecting endometrial regeneration and proliferation e.g. endometriosis and Asherman’s syndrome.

## Introduction

The endometrium is a complex tissue that cyclically regenerates every menstrual cycle in preparation for embryo implantation. Though much research has gone into understanding the endometrial mechanisms involved in the implantation event, far less is known about the tissue regeneration, akin to scarless wound healing, observed in the proliferative phase. This is important not only to understand normal endometrial physiology, but also for deciphering the pathophysiology in conditions with impaired endometrial regeneration and proliferation such as Asherman’s syndrome and endometriosis.

Stromal cells are the most abundant cell type in the endometrium and make up the mass of the regenerative endometrial *functionalis*. However, there is limited knowledge of the endometrial stromal compartment in terms of stromal-immune cell interactions, possible niches and functional subtypes. So far only one perivascular stromal cell population has been separately characterized, whereby cell surface marker expression and the transcriptomic profile were described (1, 2). This stromal subpopulation expresses CD146, platelet-derived growth factor receptor beta (PDGFRβ) and/ or sushi domain containing 2 (SUSD2) (2, 3), and has been found to exhibit some of the characteristics seen in mesenchymal stromal cells (MSCs) derived from the perivascular environment of other tissues e.g. adipose and umbilical cord isolated using CD90/ THY1, CD73 and CD105 (2, 4). Frequently these cells are considered the endometrial stromal progenitor cells(1). Apart from this population, the majority of the stromal compartment remains widely unexamined. In a recent study we sought to characterize endometrial stromal cells (eSCs) in terms of their immunomodulation of T cells and cytokine production in the proliferative phase (5). In doing so we found that all expanded eSCs express MSC surface markers thus making the distinction between fibroblasts and progenitor cells based on MSC cell surface markers problematic. eSCs were also shown to be unique in their immune interactions compared to other MSCs, especially in terms of their lack of major histocompatibility complex class II (HLA-II) expression post pro-inflammatory stimulation (5). This falls in line with previous findings describing the endometrium as a unique inflammatory milieu whereby immune cells are carefully regulated throughout the menstrual cycle (6) to promote endometrial repair instead of scarring (7).

In other tissues housing a complex stromal compartment, for example lung (8, 9), prostate (10) and lymph node (11), a diverse number of stromal subtypes with distinguishable features have been identified. Subtypes have been linked to niche and tissue location (11), as well as, specific functional properties such as response to injury by pericyte subsets (12). Given this, it is likely that the endometrial stroma holds the same level of complexity and diversity to enable the scarless repair and controlled inflammatory state of the menstrual cycle.

In the present study we set out to further investigate the endometrial stromal compartment in the proliferative phase. Using an unbiased single-cell approach and bioinformatics we aimed to draw a systematic map of endometrial stromal subsets with signature marker genes for each sub-type and propose functional characteristics.

## Results

### Unbiased single-cell analysis confirms known endometrial cell types

We set out to explore the endometrial stromal compartment in the proliferative phase to identify stromal cell complexity. After viability and quality control, isolated cells were submitted for single-cell RNA sequencing (scRNA-seq) on the 10x Genomics Chromium platform (Fig. 1A). In order to obtain an unbiased profile of the stromal compartment, cells were not *in vitro* expanded nor sorted for known stromal and progenitor markers. Transcriptomic profiles for 6,864 cells were acquired which was reduced to 6,348 after further quality control and filtering (see Materials and Methods). Using t-distributed stochastic neighborhood embedding (tSNE) to visualize the cells, seven clusters could be isolated with unique transcriptional profiles (Fig. 1B). All donors contributed equally to the cell clusters while batch effect was corrected for.

**Figure 1.**
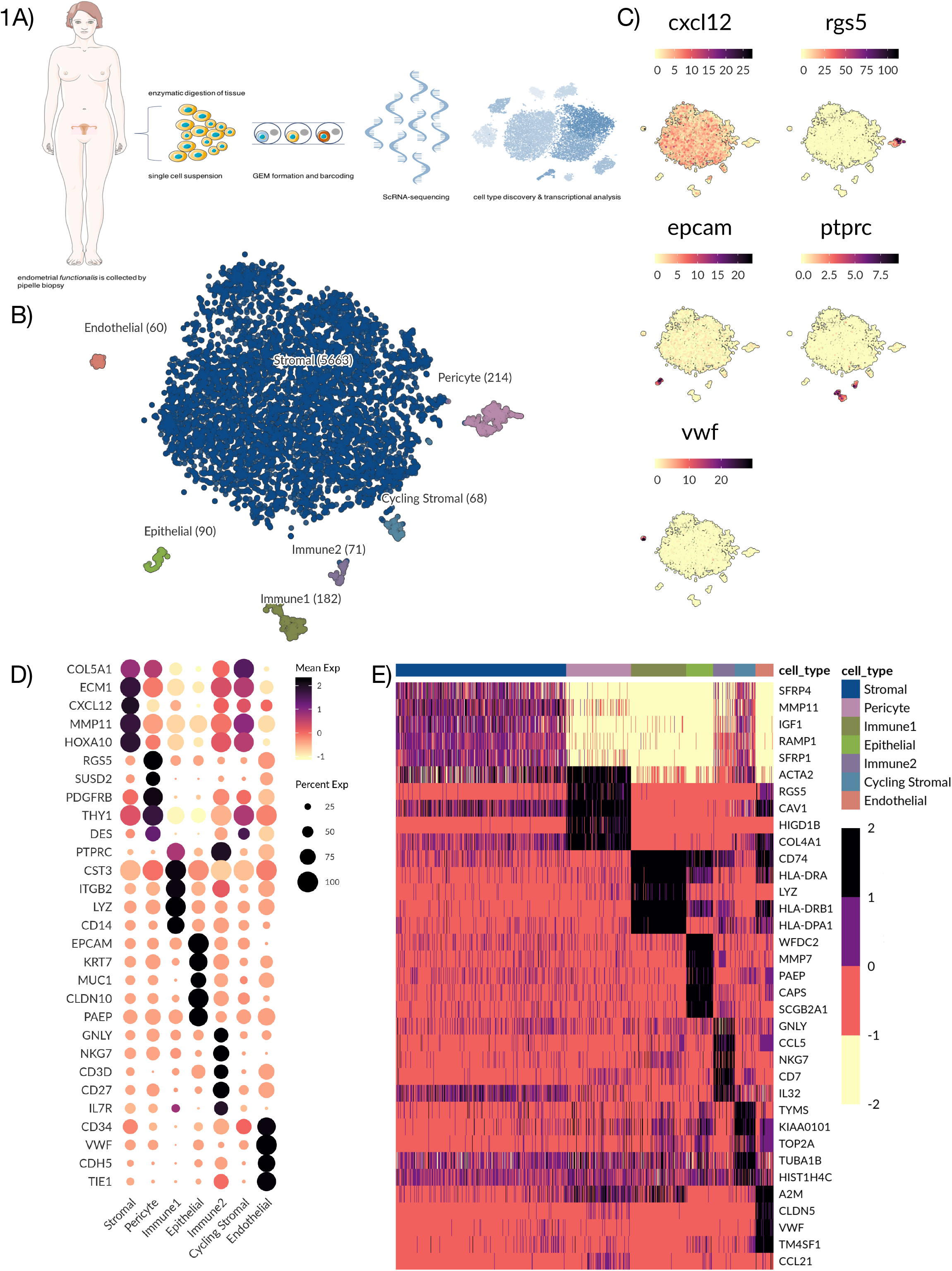
Single-cell clustering and identification of endometrial cells. **A**) Work flow of scRNA-seq and analysis of endometrial tissue. Endometrial biopsies (n=3) were collected from three healthy fertile women and enzymatically digested to a single-cell suspension. mRNA from single cells were captured and processed using the 10x genomics chromium workflow followed by Illumina sequencing. Raw reads were processed to gene counts using Cell ranger. Gene counts were then used downstream for cell type discovery and analysis. **B**) t-SNE plot visualizing clustering of scRNA-seq results from 6,348 endometrial cells. Seven clusters were detected namely epithelial cells (n=90), endothelial cells (n=60), pericytes (n=214), immune1 (n=182), immune2 (n=71), stromal cells (n=5663) and cycling stromal cells (n=68). **C**) Gene expression profiles of top marker genes identifying each cell cluster. *CXCL12* shows broad expression in stromal populations, *RGS5* expression identifies pericytes, *EPCAM* identifies epithelial cells, PTPRC identifies immune cells and *VWF* identifies endothelial cells. **D**) Dotplot showing gene expression of additional selected marker genes for each cell type to further identify clusters. Dots denote mean of normalized expression values over clusters. Immune1 displays high expression for genes indicating a monocyte phenotype, for example *CD14* and *LYZ*. Immune2 displays expression of *CD27* indicating a T cell/ NK cell phenotype. **E**) Heatmap displaying the top differentially expressed genes (rows) for each cell type (columns) based on MAST test with a minimum log fold change of 2 and adjusted p-value of 0.05. Color scale is clipped at 2.5.

In order to identify cell types, we explored the expression of different well-known marker genes in our seven clusters (Fig. 1C & D). We found unique identifier genes for each cell type: *VWF* identified endothelial cells, *EPCAM* distinguished epithelial cells and *RGS5* identified pericytes. Two clusters were identified as immune cells by expression of *PTPRC*/ CD45 (Immune1 and Immune2). The main cluster making up the bulk of the sequenced cells was identified as stromal cells with broad expression of *CXCL12* (Fig. 1C). This was to be expected as the tissue digestion protocol was optimized for eSC enrichment and eSCs make up the bulk component of the *in vivo* endometrial tissue composition. A more extensive characterization of each subset followed: with endometrial stromal progenitor and MSC markers (e.g. *SUSD2*, *PDGFRB*, *THY1*) most highly expressed in the pericyte subset (Fig. 1D). Immune1 was predominantly composed of monocytes (*CD14*) and macrophages while Immune2 included NK-cells and T-cells (*CD27*) respectively (Fig. 1D).

We applied Model-based Analysis of Single-Cell Transcriptomics test (MAST, see Materials and Methods) with a minimum log fold change of 2 and adjusted p-value of 0.05 to determine the top differentially expressed genes for all identified cell types (Fig. 1E). The stromal compartment had the least number of differentially expressed genes relative to the other subsets, suggesting a more diverse composition. Stromal signature genes were *SFRP1* and *SFRP4* involved in the Wnt-bone morphogenic protein (BMP) signaling pathway, *IGF1*, a growth factor associated with immunomodulation and regeneration, and extracellular matrix (ECM) genes e.g. *MMP11*. Pericyte signature genes were: *ACTA2*, a marker for smooth muscle actin highly expressed in the vasculature and in fibroblasts; *RGS5*, a known perivascular maker, and *CAV1*, an inhibitor of the TGFβ1 pathway regulating inflammation and ECM production. In line with this *COL4A1*, a key player in angiogenesis, was also highly expressed in the pericyte cluster.

In Immune1 there was high expression of HLA-II related genes *CD74*, *HLA-DRA*, *HLA-DR1A* and *HLA-DR1B* and bacteriolytic LYZ while in Immune2 genes relating to T cell and NK cell recruitment and activation were highly expressed (*GNLY*, *CCL5*, *NKG7*, *IL32* and *CD7*). In the epithelial subset, previously identified markers of epithelial wound repair and immunomodulation (*WFDC2*, *PAEP*, *MMP7*) were highly expressed, as well as, *CAPS* which is involved in cell-signaling and *SCGB2A1*, an androgen-regulated gene. In the endothelial subset, endothelial marker genes (*CLDN5* and *VWF*) were highly expressed along with *TM4SF1*, a marker previously associated with activated endothelial cells as well as *AM2,* which is known to limit angiogenic sprouting. A smaller stromal cluster shared the same gene expression profile as the main stromal subset but had higher levels of cell cycle related genes (*TYMS*, *KIAA0101*, *TOP2A*) as its only distinguishing feature. All of the cell type clusters we identified are known to be present in endometrial tissue and thus our results indicate that our data represents an unbiased mix of endometrial cell types.

### Heterogeneity in the endometrial stromal compartment and possible stromal subtypes revealed by single-cell analysis

To further characterize the stromal compartment, stromal cells were subset from the main dataset for additional in-depth analysis. As the cycling stromal cells had a different cell cycle phenotype they were excluded from downstream analysis. Using Uniform Manifold Approximation and Projection (UMAP) analysis the stromal cells were re-clustered, forming ten independent populations (Fig. 2A). UMAP analysis uses spatial placement to illustrate the association of different clusters to one another. Although perivascular cells are not considered stromal cells per se, their role in stromal regeneration is established and as such they were included in the initial clustering to determine how and to which stromal clusters they relate (Fig. 2A).

**Figure 2.**
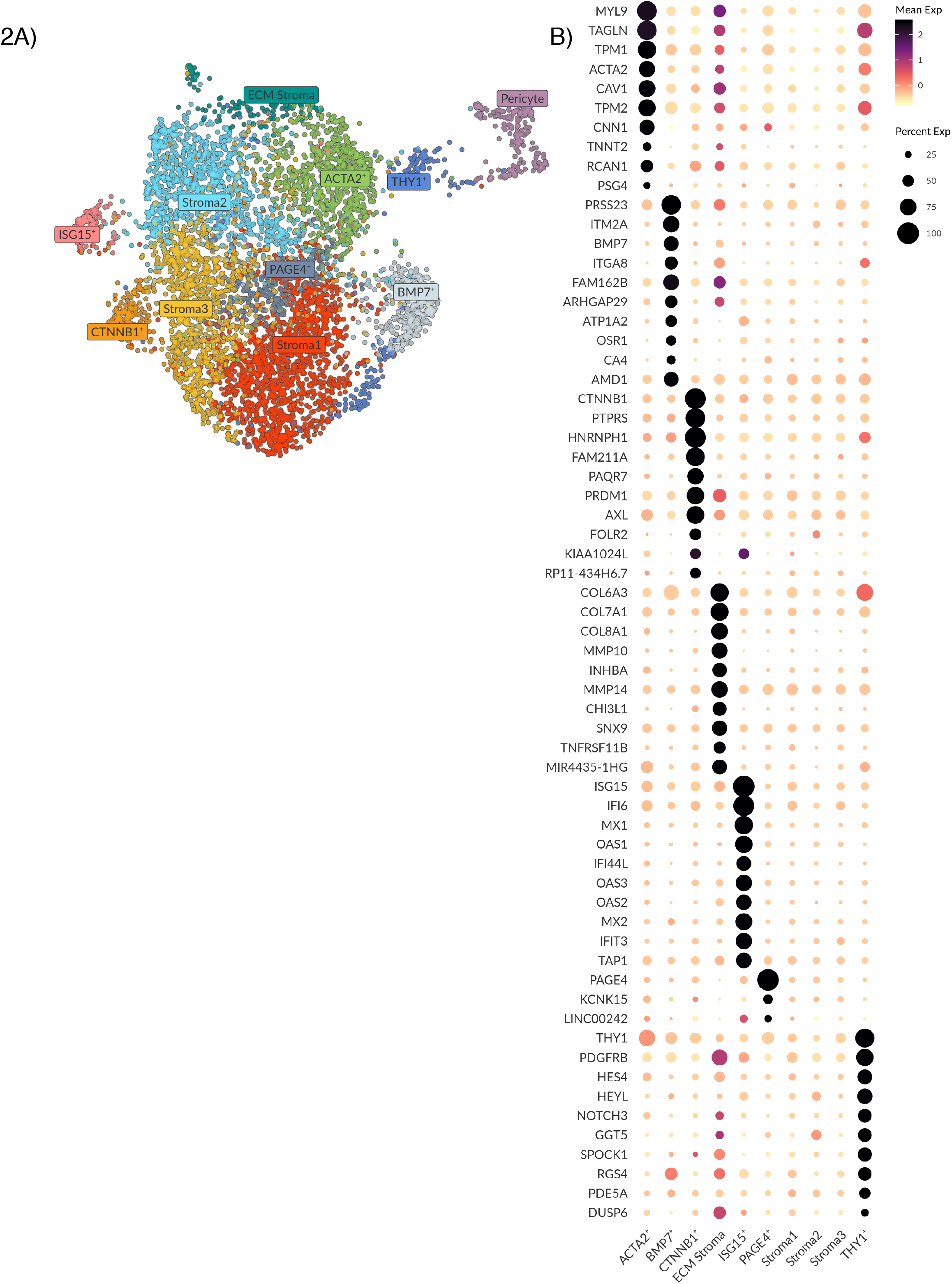
Analysis and subtyping of the endometrial stromal cell compartment. **A**) UMAP plot showing ten clusters of endometrial stromal cells and a cluster of pericytes on the far right. Clusters are labelled as per their identified expression profile in B. **B**) Dotplot showing the top differentially expressed genes (rows) for the ten stromal clusters (columns). Genes were selected based on MAST test with a minimum log fold change of 2 and adjusted p-value of 0.05. Note that the color scale is clipped at 2.5. Stroma 1, 2 and 3 do not show any unique expression. The PAGE4+ cluster shows a very biased differential expression for *PAGE4* only. The ECM, ACTA+ and BMP7+ clusters show a higher expression for genes involved in ECM breakdown, remodeling and organization. The CTNNB1+ cluster show higher expression of genes involved in epithelial regulation and innate immunity. The ISG15+ cluster show higher expression for genes involved in innate immunity. The THY1+ cluster shows higher expression of genes involved in Notch signaling.

Using the MAST test with a minimum log fold change of 2 and adjusted p-value of 0.05, the top ten distinguishing genes for each stromal subset were determined to generate a profile (Fig. 2B). Three clusters (Stroma 1, 2 and 3) did not exhibit any unique expression profiles compared to the other clusters. We interpret this as these cells being the baseline stromal cells that comprise the bulk of endometrial stroma. Closely related to these clusters we saw one small cluster with biased expression for *PAGE4* (Fig. 2B). Based on our extensive literature search and the limited number of genes distinguishing this subset, the PAGE4+ subset was not further explored.

In the remaining six stromal populations the gene profiles were characterized and labelled based on significant genes (Fig. 2B). The ACTA2+ population had a gene signature (*MYL9*, *TAGLN*, *TPM1*, *ACTA2*, *TPM2*, *TNNT2* and *CNN1*) suggestive of activated fibroblasts and smooth muscle (13, 14), specifically genes modulating actin and myosin interactions. Adjacent to this cluster, the ECM population had the highest expression of collagens and matrix metalloproteinases (MMPs) genes (*COL6A3*, *COL7A1*, *COL8A1*, *MMP10* and *MMP14*), which are components of the basement membrane (e.g. surrounding vasculature) and vital in the regulation of tissue remodeling and homeostasis. The BMP7+ population had genes involved in myofibroblast differentiation, epithelial mesenchymal transition (EMT) and in TGFβ1-WNT signaling (*PRSS23*, *ITGA8*, *BMP7*, *ITM2A* and *ARHGAP29*) (15–18). We hypothesize these three stromal clusters (ACTA+, ECM and BMP7+) represent stromal subtypes active in ECM breakdown, remodeling and organization which are important processes during proliferation, tissue repair and regeneration occurring in the endometrium, during, and after menstruation.

In one stromal cluster we noted a high expression of *CTNNB1* (Fig. 2B). This gene has been previously linked to Wnt signaling and stromal cell regulation of epithelial proliferation and differentiation in wound healing (19, 20). Other genes in this population have also been associated with epithelial-stromal interactions, as well as, innate immune responses, specifically M2 anti-inflammatory macrophage polarization (*PTPRS*, *PAQRF*, *PRDM1*, *AXL* and *FOLR2*) (21–24). Similarly, in the ISG15+ population, most genes are involved in interferon signaling and innate immunity functions (*ISG15, IF16, MX1, OAS1, IF144L, OAS3, OAS2, MX2, IFIT3*) (25, 26), suggestive of an activated stromal cell state/ a subtype active in innate immune immunomodulation.

Lastly, the THY1+ stromal perivascular population was distinguished by high *THY1* expression, which is a surface glycoprotein commonly used as an MSC marker. This cluster also had the highest gene expression associated with Notch signaling (*HES4, HEYL, NOTCH3*), a pathway involved in cell-cell signaling, cell plasticity and differentiation, as well as, genes associated with the pericyte (*PDGFRB, PDE5A*) (27, 28).

### Single-cell analysis of endometrial pericyte cells reveals two distinct subtypes

Using the known pericyte marker RGS5 we identified one cell population containing pericytes (Fig. 3A) (28). As pericytes have previously been linked to endometrial stromal regeneration we wanted to further explore this cell subset (2, 29). Isolating pericytes from the main dataset and applying UMAP analysis to determine differential gene expression and higher complexity in this environment yielded two separate populations (Pericyte1 and Pericyte2) (Fig. 3B).

**Figure 3.**
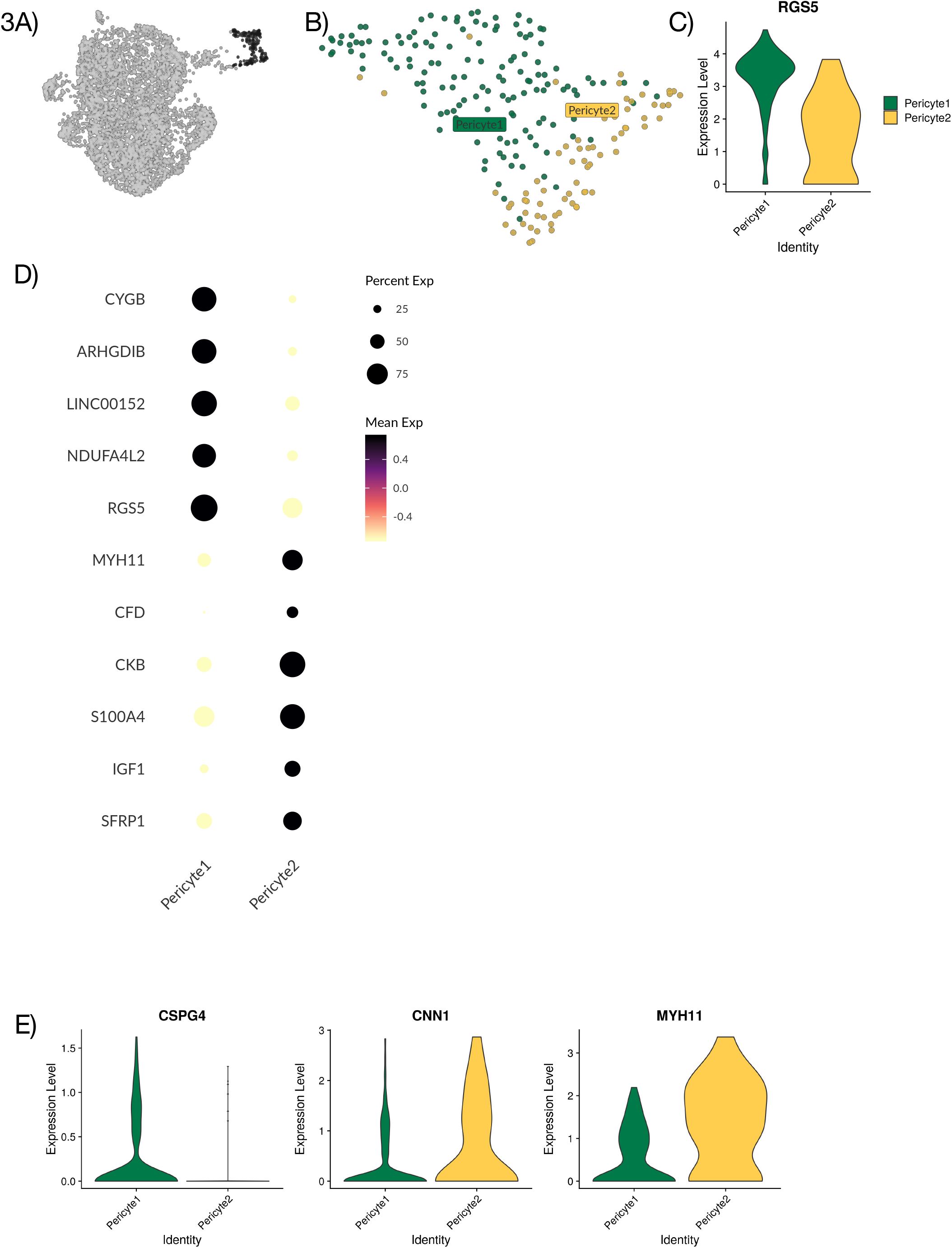
Analysis and subtyping of endometrial pericytes. **A**) UMAP plot of scRNA-seq results of endometrial stromal cells and pericytes highlighting cells (in black) expressing pericyte marker *RGS5*. These cells were subset for further analysis. **B**) UMAP plot of scRNA-seq results visualizing clustering of endometrial pericytes into two separate clusters, Pericyte1 and Pericyte2. **C**) Violin plot displaying full distribution of *RGS5* expression in Pericyte1 and Pericyte2. **D**) Dotplot showing mean expression of top differentially expressed genes for Pericyte1 and Pericyte2 after MAST test with minimum log fold change of 1.5 and adjusted p-value of 0.05. **E**) Violin plots showing full distribution of marker genes *CSG4, CNN1* and *MYH11* in Pericyte1 and Pericyte2 with the profile *CSPG4*^+^ *CNN1*^*low*^ *MYH11*^*low*^ for Pericyte1 and *CSPG*^−^ *CNN1*^*high*^ *MYH11*^*high*^ for Pericyte2.

In order to explore the identities of Pericyte1 and Pericyte2, the expression levels of the perivascular marker RGS5 were compared, revealing that Pericyte1 had a higher mean expression than Pericyte2 (Fig. 3C). By applying MAST analysis with a minimum log fold change of 1.5 and adjusted p-value of 0.05, the differentially regulated genes between the two populations were determined, with *CYGB, ARGHDIB, LINC00152* and *NDUFA4L2* specific to Pericyte1 and *MYH11*, *CKB, CFD, S100A4, IGF1* and *SFRP1* specific to Pericyte2 (Fig. 3D). *MYH11* has previously been used as a marker for mature smooth muscle cells (SMCs) in the perivascular niche (30). To further investigate if Pericyte2 could represent a SMC like pericyte subtype, the expression of additional reported markers *CSPG4*, *CNN1* and *MYH11* in both Pericyte1 and Pericyte2 was validated (30) (Fig. 3E). Pericyte1 presented a *CSPG4*^+^ *CNN1*^*low*^ *MYH11*^*low*^ profile and Pericyte2 had a *CSPG*^−^ *CNN1*^*high*^ *MYH11*^*high*^ profile. This is consistent with Pericyte1 representing a classic pericyte/ mural cell population, and Pericyte2 representing a SMC-like or contractile pericyte.

### Transcriptional expression of PDGFRB, MCAM, SUSD2 and THY1 extends across the greater perivascular niche

Endometrial stromal regeneration has been said to be orchestrated by stromal progenitor cells in the perivascular environment, as such we attempted to establish a hierarchy between cell populations (ACTA2+, THY1+, Pericyte1 and Pericyte2) presenting gene profiles indicative of a perivascular and smooth muscle actin identity, thereby establishing more specific marker genes (Fig. 4A). *THY1* expression was seen to increase in the stromal subsets, with increased proximity to the pericyte as per UMAP analysis, where local and global structures are preserved and thus distance between subsets indicates similarity of subsets (Fig. 4B). The four populations were subset and integrated, UMAP analysis was performed (Fig. 4C) and relative scaled expression across the subset of *PDGFRB*, *MCAM*, *SUSD2* and *THY1* was determined (Fig. 4D). *PDGFRB* was highly expressed in Pericyte1 and THY1+ cells. *MCAM* and *SUSD2* were highly expressed in Pericyte1 and Pericyte 2. *THY1* was highly expressed in Pericyte1 and THY1+ cells. No specific marker distinguished the THY1+ cells from other cell populations in the subset (Fig. 4D). We applied RNA velocity (Velocyto, see Materials and Methods) analysis on the subset to determine developmental trajectory (Fig. 4E). Within each cell on the principal component analysis (PCA) plot, amplitude and direction of the arrow provides information on developmental trajectory. Pericyte1 and Pericyte2 showed long active arrows with opposing directional progression to different states. Within the THY1+ cells there were two sub-populations: one closer to the ACTA2+ cells, with short arrows suggesting a steady state, while the other subset had longer arrows committing towards Pericyte1. The ACTA2+ cells presented a uniform identity with the shortest arrows overall, possibly an example of a more committed state. Their trajectory was in the opposite direction of Pericyte1 and Pericyte2. Finally, we sought to identify more specific marker genes for the different populations within the perivascular environment so they can be more easily distinguished relative to the other populations in this environment by applying MAST analysis with a minimum log fold change of 1.5 and adjusted p-value of 0.05 (Fig. 4F). *TXN, KRT19, TGFBI, VCAN* and *GLIPR1* specifically identified the ACTA2+ cells. *HES1*, *SPOCK1*, *HTRA3*, *CHST1* and *IGFBP3* identified THY1+ cells. *ARHGDIB, NDUFA4L2, RGS5, CYGB* and *ANGPT2* identified Pericyte1. *FXUD1*, *SOD3*, *SLIT3*, *LG14* and *ACTG2* identified Pericyte2. Overall, these gene profiles provide more specific marker genes within the perivascular environment as current gene profiles appear to be more telling of cell location rather than cell type.

**Figure 4.**
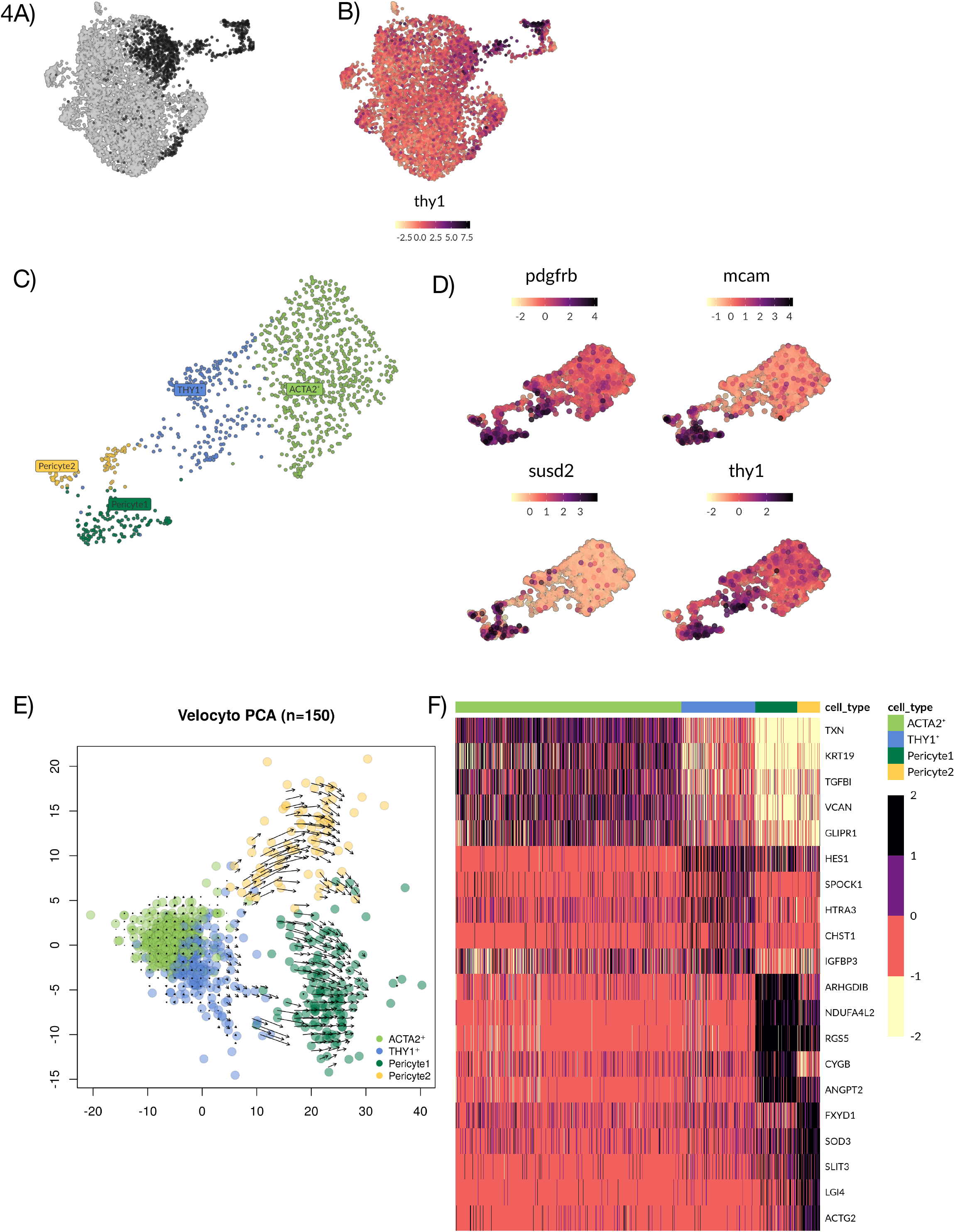
Analysis of subsets ACTA2+, THY1+, Pericyte1 and Pericyte2 clusters with. **A**) UMAP plot of endometrial stromal cells and pericytes highlighting cell types ACTA2+, THY1+ and pericyte 1 and 2 (in black). **B**) UMAP plot of endometrial stromal cells and pericytes showing increased expression of *THY1* with increasing proximity to the pericyte cluster at the top right. **C**) UMAP plot showing groups ACTA2+, THY1+ and Pericyte 1 and 2. **D**) UMAP plot showing expression pattern of genes *PDGFRB*, *MCAM*, *SUSD2* and *THY1.* None of these markers can exclusively identify any of the clusters mentioned in C. **E**) PCA scatterplot showing RNA velocity. Predicted developmental trajectory between clusters is displayed as a vector field. Short arrows indicate a steady state and long arrows indicate active progression towards a differentiated state. Cells differentiate along the direction of the arrow, here indicating some of the THY1+ cells committing towards the Pericyte1. **F**) Heatmap showing the top differentially expressed genes (rows) for each cell cluster (columns) based on MAST test with minimum log fold change of 1.5 and adjusted p-value of 0.05.

### Validation of endometrial stromal and pericyte subtypes in an external data set reveals distinguishable subtypes persisting throughout menstrual cycle

In order to validate our findings of endometrial stromal and pericyte sub-types we imported external human scRNA-seq data from the maternal-fetal interface by Vento-Tormo *et al*., (2018) (31). The dataset was filtered for the maternal CD45^−^ decidual fraction (12,544 cells) providing a suitable comparison to the endometrial stromal compartment and perivascular cells. Maternal decidua is endometrium following decidualization and placentation. As part of the original analysis three decidual stromal populations had been explored (dS1, dS2 and dS3) and two perivascular populations (dP1 and dP2) (Fig. 5A). Similar workflow as initially described was applied to the subset data of interest. Louvain clustering was used in Seurat to identify 10 clusters (Fig. 5B). In order to investigate if any of these clusters represented the subtypes that had been identified in our data set, we searched for the expression of some of our distinguishing marker genes in the clusters (Fig. 5C). Gene expression profiles identified one cluster as the CTNNB1+ population (Fig. 5Ci), another as the ACTA2+ population (Fig. 5Cii), and a third as the ISG15+ stromal subtype (Fig. 5Ciii). This suggests that these subtypes/ cell states are persistent players in the endometrial stromal compartment throughout the menstrual cycle and in early pregnancy. In order to investigate our pericyte subtypes, cells were selected for *RGS5* expression in the external dataset (Fig. 5D). Using the markers *CYGB*, *ARGHDIB* and *NDUFA4L2* we could readily identify one cell cluster as corresponding to Pericyte1 and using *MYH11* expression Perictye2 was identified (Fig. 5E).

**Figure 5.**
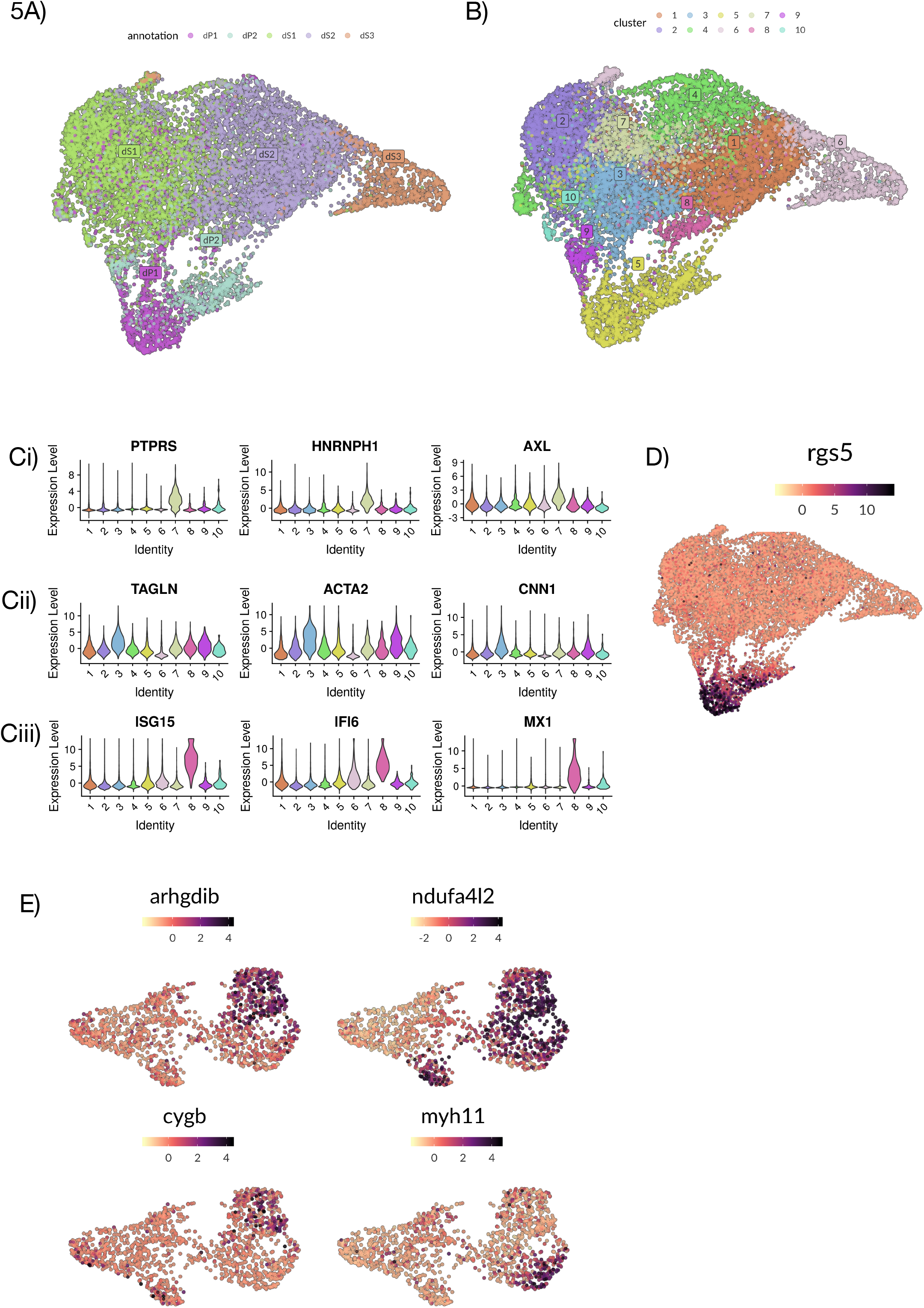
Analysis and identification of endometrial stromal and pericyte subtypes in the maternal-fetal interface. **A)** UMAP plot of scRNA-seq data of the maternal CD45^−^ decidual fraction from the maternal-fetal interface. Annotation of cell clusters according to original publication by Vento-Tormo *et al.* (31) which identifies which identifies pericytes (dP1 and dP2) and stromal cells (dS1, dS2 and dS3). **B)** UMAP plot of scRNA-seq data of the maternal CD45^−^ decidual fraction from the maternal-fetal interface displaying ten clusters using the same parameters applied in our dataset. **C**) Violin plots showing full distribution of marker gene expression for each population. i) CTNNB1+ epithelial regulator; *PTPRS*, *HNRNPH1* and *AXL* ii) ACTA2+ activated fibroblast; *TAGNLN*, *ACTA2* and *CNN1* iii) ISG15+ innate immunity stromal subtype; *ISG15*, *IFI6* and *MX1.* **D**) Scatter plot showing RGS5 expressing cells in the UMAP projection. **E**) Scatter plot of selected RGS5+ cells and their scaled gene expression of *CYGB*, *ARGHDIB*, *NDUFA4L2* and *MYH11* identifying the top right cells as Pericyte1 and the bottom right cells as Pericyte2.

## Discussion

Understanding healing and growth mechanisms is important when explaining deregulation in benign gynecological disorders which present with inflammatory/ proliferative imbalance (32, 33). Furthermore, the endometrium’s unique regenerative mechanism could provide us with knowledge to develop cell and gene therapy in order to combat chronic diseases. Using unbiased single-cell transcriptional data, this study indicates that the nature of the endometrial stromal compartment is more complex than previously assumed. Undeniably, the endometrial perivascular environment holds many unanswered questions about its role as a progenitor cell niche and central regulator of endometrial regeneration. However, equally important are the other niches which coordinate controlled stromal proliferation, the ECM’s ever-changing composition and inflammatory homeostasis. The six gene expression signatures of stromal subsets we have described in this work; of which three are retained throughout decidualization and early pregnancy, providing a starting point towards understanding the complexity of the greater stromal compartment. At this point, further follow up experiments are required to say whether transcriptomic heterogeneity in a cell population may reveal previously unknown cell types, cell states or cell niches *in vivo*. Thus, all our stromal subsets warrant translational validation and further exploration to provide functional meaning to the transcriptional profiles.

Previously, we observed that eSCs have a unique immunomodulatory phenotype, describing a lack of HLA II cell surface expression in pro-inflammatory conditions (IFNγ and TNF-α licensing) in contrast to other stromal cells. Furthermore, they limit inflammation by reducing CD4+ T helper cells proliferation while shifting them towards an effector memory phenotype (5). In the current study, particularly the ISG15+ population revealed an interferon regulated gene expression profile suggestive of an activated stromal population, as seen in inflammatory settings such as tissue regeneration following menstruation (34). Similarly, the CTNNBI+ population included genes linked to M2 macrophage polarization e.g. *AXL*, *FOLR2* (22–24). M2 macrophages are involved in remodeling tissue and secreting anti-inflammatory cytokines, with evidence suggesting stromal cells may be able to polarize macrophages to a M2 state via paracrine mechanisms (35, 36). Immunomodulation studies *in vitro* could provide us with better understanding of the ISG15+ population and the CTNNBI+ population. While computational tools like CellPhone DB with extensive endometrial immune cell sequencing data could provide further information about specific eSC mechanisms in tissue regeneration and the immunomodulation of specific lymphocytes (37).

The ACTA2+ subset clearly distinguished itself from other stromal subsets by its high expression of smooth muscle actin and myosin regulating genes. Further delineation of the population’s identity was complicated by the inability to discriminate between a perivascular smooth muscle location, an activated fibroblast or terminally differentiated myofibroblast based on a transcriptional profile alone. *ACTA2* has historically been used as a marker in both settings (13, 14). Based on the RNA-velocity analysis, ACTA2+ cells presented a steady state with the lowest differentiation capacity relative to other cell populations in the perivascular environment, aligning their identity more with an activated fibroblast/ myofibroblast. ACTA2+ cells were also highlighted in the analysis by Vento-Tormo *et al*. with gene and protein expression observed in the perivascular environment, as well as, the decidua spongiosa adjacent to the myometrium (31). Thus, there is a general need to find specific markers to discriminate between smooth-muscle proximity and myofibroblast activation to better determine each of these cells’ role in regeneration (38).

The endometrial pericyte appears to be made up of two subsets, with Pericyte1 potentially representing a more classical pericyte/ mural cell while Pericyte2 revealing a gene profile more aligned with vascular SMCs as per the gene profiles introduced by Kumar *et al.* (30). However, the greater pericyte field has no specific marker profile which indisputably and regardless of developmental stage identifies pericytes from vascular SMCs or MSCs for that matter (38, 39). Common markers include *ACTA2*, *PGDFRB*, *DES*, *RGS5* and *CSPG4* (38). This leads to the question of where the perivascular cells end and the stromal compartment with myofibroblasts and progenitors begins. PDGFRβ, MCAM (2) and SUSD2 (3) have previously been used as markers in the endometrium to identify stromal progenitors, however our data suggests these genes are expressed by all cells in the perivascular environment, with the highest gene expression in the Pericyte1 and Pericyte2. Further investigation as to whether these markers are more discriminatory at a protein level is needed, especially as stromal progenitors and pericytes are isolated and enriched using similar cell-surface markers. THY1/ CD90 in combination with CD73 and CD105 are used as MSC markers to identify endometrial stromal progenitors (40). Although our data shows differential expression of THY1 across the stromal compartment and pericyte, it is not exclusive to a progenitor cell or the pericyte. THY1 is expressed on the cell surface of a number of cells, including fibroblasts (5, 41), and while it is associated with undifferentiated states in stromal and hematopoietic cells, it is a marker progressive maturation in neurons (42). This demonstrates the complexity of its biological role beyond being a marker and the importance of the cellular microenvironment (42). Overall, our findings suggest that existing endometrial markers for progenitor cells in the perivascular environment are not specific enough on a gene level to distinguish between the key cells in this niche. This makes it difficult to distinguish them from one another and determine their individual role in regeneration.

A possible reason why multiple stromal populations could not be validated in the external dataset was due to the considerable effect of decidualization on stromal cell morphology and their immunophenotype. Naturally tissue dissociation will also alter the transcriptome possibly distinguishing the two datasets (43).

Our scRNA-seq study has provided descriptive data on numerous stromal subsets and exploratory routes to investigate their role in endometrial regeneration. Multiple stromal environments exist, which include different cell types, cell states, ECM compositions and immune settings. Furthermore, this study provides information about the complexity of the perivascular environment, revealing how very few genes are truly unique to one cell population, making it complex to distinguish between progenitor cells, fibroblasts, smooth muscle cells and mural cells. This is particularly the case, if gene and protein expression are heavily affected by microenvironment rather than a specific cell type.

## Materials and Methods

### Healthy Endometrial Donor Material

Endometrial proliferative stage *functionalis* samples were obtained from healthy volunteers (n = 3), aged 30–32 years. The study was approved by the regional ethics committee of Karolinska Institutet, Stockholm, Sweden (ethical approval reference numbers DNR: 2015/367-31/4 and 2017/216-31). Written informed consent was obtained from all participating women. All donors had normal menstrual cycles (25–35 days) and were proven fertile (at least one confirmed pregnancy). Women were examined for the absence of hormonal diseases, uterine pathologies (e.g., endometriosis, polycystic ovary syndrome and/or previous infertility records), and sexually transmitted diseases (HIV, Chlamydia trachomatis-DNA, and Gonococci-DNA). None of the women had used hormonal contraception or an intrauterine device for a minimum of 3 months prior to biopsy. Biopsies were obtained without cervical dilation or local anesthesia, using a pipelle aspirator from Cooper Surgical (Trumbull, USA).

### Endometrial Single Cell Isolation

Once the endometrial biopsies were collected, they were stored in MACS® Tissue Storage Solution (Miltenyi Biotec, Lund, Sweden) at 4°C until further processing. The sample was washed with phosphate-buffered saline (PBS; Sigma-Aldrich, Taufkirchen, Germany) and minced into 2mm^3^ pieces. The tissue was gently digested in a filter sterilized Dispase II solution (0.5 U/ml; Sigma Aldrich, Taufkirchen, Germany) in complete media composed of DMEM-F12 (Thermo Fisher Scientific, Dreieich, Germany) with 10% (v/v) fetal calf serum (FCS; Thermo Fisher Scientific) at 4°C overnight. The tissue solution was manually disaggregated, washed with complete media and centrifuged at 200×g for five minutes. The tissue was further digested with filter sterilized Collagenase III (150U/ml; Worthington, Lakewood, USA) and DNAse (139U/ml; Sigma Aldrich) in complete media with agitation for 45 min at 37 °C. Once the tissue was completely dissociated, the cells were washed in complete media and centrifuged at 200xg for five minutes. The cell pellet was treated with 1 ml of Red Blood Cell Lysis Buffer (Roche, Solna, Sweden) for five minutes at room temperature. The reaction was stopped with the addition of complete media, and the cells washed and centrifuged at 200×g for five minutes. Cells were counted and viability assessed using the TC20™Automated Cell Counter (BioRad, Gothenburg, Sweden). The cells were resuspended in PBS containing bovine serum albumin (400μg /ml; Sigma Aldrich) at a cell concentration of 1000 cells/μl.

### Single-cell library preparation and mRNA sequencing (scRNA-seq) of proliferative phase endometrial cells with 10x Genomics

The endometrial cell suspensions were delivered to the Eukaryotic Single Cell Genomics Facility (ESCG, SciLifeLab, Stockholm, Sweden), prepared and loaded on a 10x Genomics Chromium Controller instrument for single-cell gel bead-in-emulsion (GEM) formation and barcoding using the kit Chromium Single Cell 3’ Gel Bead Kit v2. GEM reverse transcription was performed. Once cDNA was generated, amplified by PCR and cleaned, the sequencing libraries were constructed according to the manufacturer’s instructions using the Chromium Single Cell 3’ Library Kit v2.

Three runs of scRNA-seq of uncultured endometrial cells were performed with one run per sample (n=3). Each run consisted of one sample/sequencing lane using the Illumina 2500 instrument. Approximately 3,000 cells were sequenced per sample with a sequencing depth of 50,000 reads per sample.

### ScRNA-seq data analysis

ScRNA-seq output files were converted with Cell Ranger 2.1.1 and aligned to the hg19 transcriptome using STAR mapper (44). Cell Ranger was used to process raw sequencing data. Analysis of filtered cells was performed in R versions 3.6.0 and 3.6.1 using Seurat suite versions 3.1.2 and 3.1.3 (45, 46). Initial quality control measures were performed to exclude potential doublets and dying cells, cells expressing 200 – 5,000 genes, and less than 10% of mitochondrial genes were kept, resulting in 6,348 cells in total. ERCC, RPL and RPS genes were removed. All cells were normalized according to their cell cycle stage and the data was corrected for batch effect using the integration tool sctransform (47). The dimensionality of the data was reduced using PCA. Elbow plot was used to selected top PCs which were used downstream for Louvain clustering and visualization using tSNE and UMAP. “Reference-Based Single-Cell RNA-Seq Annotation” tool SingleR (48) was run to broadly identify cell types. Final cell type labels were established after manual assessment using known marker genes. Once cells were identified, the marker genes for each subset were determined by differential gene expression tests using MAST (49). Larger groups of cells were further subset with downstream analysis using the same Seurat steps again. Velocyto was used to evaluate cell lineage by cell dynamics and RNA velocity (50).

An external dataset was used to validate gene profiles found in the endometrial dataset. This dataset was first published by Vento-Tormo *et al*., 2018 (VT)(31). The data were downloaded at: https://www.ebi.ac.uk/arrayexpress/experiments/E-MTAB-6701/. The VT dataset was integrated with our data to investigate similarity in cell types and expression profiles. See the original publication for further details on the VT data and their analyses.

### Statistics

Changes in scaled gene expression between cell types were calculated using the MAST test (49). MAST uses a generalized linear model framework that considers the bimodal nature of single-cell expression data due to stochastic drop-outs. MAST offers a differential gene expression test that is custom tailored for single-cell count data. A log fold change of 2 or 1.5 and adjusted p-value of 0.05 was applied to determine highly unique genes in different subsets. The adjusted p-value/ false discover rate was based on Bonferroni correction using all genes in the dataset.

## Abbreviations

ECM: extracellular matrix
EMT: epithelial mesenchymal transition
eSC: endometrial stromal cell
FBS: fetal bovine serum
GEM: single-cell gel bead-in-emulsion
MAST: Model-based Analysis of Single-Cell Transcriptomics
MMP: matrix metalloproteinase
MSC: mesenchymal stromal cell
PBS: phosphate-buffered saline
PCA: principal component analysis
scRNA-seq: single-cell RNA sequencing
SMC: smooth muscle cell
tSNE: t-distributed stochastic neighborhood embedding
UMAP: uniform manifold approximation and projection

## Acknowledgments

The authors would like to acknowledge the WHO collaborating center for research and research training in human reproduction at Karolinska Institutet and Karolinska University Hospital for helping in collecting biopsies, and the healthy women who participated in this study. Funding was received from Jane and Dan Olsson Foundation, The Swedish research council (2017-00932) and Karolinska Institutet (KID) (2-3591/2014). Furthermore, this project received bioinformatic support from NBIS (National Bioinformatics Infrastructure Sweden).

## Notes

### Competing Interest Statement

The authors have declared no competing interest.

### Summary of Updates

Added acknowledgments section.

## References

1. Chan RW, Schwab KE, Gargett CE. Clonogenicity of human endometrial epithelial and stromal cells. Biol Reprod. 2004;70(6):1738–50.

2. Schwab KE, Gargett CE. Co-expression of two perivascular cell markers isolates mesenchymal stem-like cells from human endometrium. Human reproduction (Oxford, England). 2007;22(11):2903–11.

3. Masuda H, Anwar SS, Buhring HJ, Rao JR, Gargett CE. A novel marker of human endometrial mesenchymal stem-like cells. Cell transplantation. 2012;21(10):2201–14.

4. Dominici M, Le Blanc K, Mueller I, Slaper-Cortenbach I, Marini F, Krause D, et al. Minimal criteria for defining multipotent mesenchymal stromal cells. The International Society for Cellular Therapy position statement. Cytotherapy. 2006;8(4):315–7.

5. Queckbörner S, Syk Lundberg E, Gemzell-Danielsson K, Davies LC. Endometrial stromal cells exhibit a distinct phenotypic and immunomodulatory profile. Stem cell research & therapy. 2020;11(1):15.

6. Bonatz G, Hansmann ML, Buchholz F, Mettler L, Radzun HJ, Semm K. Macrophage- and lymphocyte-subtypes in the endometrium during different phases of the ovarian cycle. Int J Gynaecol Obstet. 1992;37(1):29–36.

7. Kaitu’u-Lino TJ, Morison NB, Salamonsen LA. Neutrophil depletion retards endometrial repair in a mouse model. Cell Tissue Res. 2007;328(1):197–206.

8. Du Y, Guo M, Whitsett JA, Xu Y. ‘LungGENS’: a web-based tool for mapping single-cell gene expression in the developing lung. Thorax. 2015;70(11):1092–4.

9. Xie T, Wang Y, Deng N, Huang G, Taghavifar F, Geng Y, et al. Single-Cell Deconvolution of Fibroblast Heterogeneity in Mouse Pulmonary Fibrosis. Cell Rep. 2018;22(13):3625–40.

10. Henry GH, Malewska A, Joseph DB, Malladi VS, Lee J, Torrealba J, et al. A Cellular Anatomy of the Normal Adult Human Prostate and Prostatic Urethra. Cell Rep. 2018;25(12):3530–42.e5.

11. Rodda LB, Lu E, Bennett ML, Sokol CL, Wang X, Luther SA, et al. Single-Cell RNA Sequencing of Lymph Node Stromal Cells Reveals Niche-Associated Heterogeneity. Immunity. 2018;48(5):1014–28.e6.

12. Goritz C, Dias DO, Tomilin N, Barbacid M, Shupliakov O, Frisen J. A pericyte origin of spinal cord scar tissue. Science. 2011;333(6039):238–42.

13. Liguori TTA, Liguori GR, Moreira LFP, Harmsen MC. Fibroblast growth factor-2, but not the adipose tissue-derived stromal cells secretome, inhibits TGF-β1-induced differentiation of human cardiac fibroblasts into myofibroblasts. Scientific reports. 2018;8(1):16633.

14. Liu R, Jin JP. Calponin isoforms CNN1, CNN2 and CNN3: Regulators for actin cytoskeleton functions in smooth muscle and non-muscle cells. Gene. 2016;585(1):143–53.

15. Chen IH, Wang HH, Hsieh YS, Huang WC, Yeh HI, Chuang YJ. PRSS23 is essential for the Snail-dependent endothelial-to-mesenchymal transition during valvulogenesis in zebrafish. Cardiovasc Res. 2013;97(3):443–53.

16. Matsushima S, Aoshima Y, Akamatsu T, Enomoto Y, Meguro S, Kosugi I, et al. CD248 and integrin alpha-8 are candidate markers for differentiating lung fibroblast subtypes. BMC Pulm Med. 2020;20(1):21.

17. Leinhos L, Peters J, Krull S, Helbig L, Vogler M, Levay M, et al. Hypoxia suppresses myofibroblast differentiation by changing RhoA activity. J Cell Sci. 2019;132(5).

18. Boeuf S, Börger M, Hennig T, Winter A, Kasten P, Richter W. Enhanced ITM2A expression inhibits chondrogenic differentiation of mesenchymal stem cells. Differentiation. 2009;78(2-3):108–15.

19. Miyazaki K, Dyson MT, Coon VJ, Furukawa Y, Yilmaz BD, Maruyama T, et al. Generation of Progesterone-Responsive Endometrial Stromal Fibroblasts from Human Induced Pluripotent Stem Cells: Role of the WNT/CTNNB1 Pathway. Stem Cell Reports. 2018;11(5):1136–55.

20. Stewart CA, Wang Y, Bonilla-Claudio M, Martin JF, Gonzalez G, Taketo MM, et al. CTNNB1 in mesenchyme regulates epithelial cell differentiation during Mullerian duct and postnatal uterine development. Mol Endocrinol. 2013;27(9):1442–54.

21. Bunin A, Sisirak V, Ghosh Hiyaa S, Grajkowska Lucja T, Hou ZE, Miron M, et al. Protein Tyrosine Phosphatase PTPRS Is an Inhibitory Receptor on Human and Murine Plasmacytoid Dendritic Cells. Immunity. 2015;43(2):277–88.

22. Kanzaki R, Naito H, Kise K, Takara K, Eino D, Minami M, et al. Gas6 derived from cancer-associated fibroblasts promotes migration of Axl-expressing lung cancer cells during chemotherapy. Scientific reports. 2017;7(1):10613.

23. Myers KV, Amend SR, Pienta KJ. Targeting Tyro3, Axl and MerTK (TAM receptors): implications for macrophages in the tumor microenvironment. Mol Cancer. 2019;18(1):94-.

24. Puig-Kröger A, Sierra-Filardi E, Domínguez-Soto A, Samaniego R, Corcuera MT, Gómez-Aguado F, et al. Folate receptor beta is expressed by tumor-associated macrophages and constitutes a marker for M2 anti-inflammatory/regulatory macrophages. Cancer research. 2009;69(24):9395–403.

25. Wichit S, Hamel R, Zanzoni A, Diop F, Cribier A, Talignani L, et al. SAMHD1 Enhances Chikungunya and Zika Virus Replication in Human Skin Fibroblasts. Int J Mol Sci. 2019;20(7).

26. Patel MV, Shen Z, Rossoll RM, Wira CR. Estradiol-regulated innate antiviral responses of human endometrial stromal fibroblasts. American journal of reproductive immunology (New York, NY: 1989). 2018;80(5):e13042.

27. Liu H, Kennard S, Lilly B. NOTCH3 expression is induced in mural cells through an autoregulatory loop that requires endothelial-expressed JAGGED1. Circ Res. 2009;104(4):466–75.

28. Bondjers C, Kalén M, Hellström M, Scheidl SJ, Abramsson A, Renner O, et al. Transcription profiling of platelet-derived growth factor-B-deficient mouse embryos identifies RGS5 as a novel marker for pericytes and vascular smooth muscle cells. The American journal of pathology. 2003;162(3):721–9.

29. Spitzer TL, Rojas A, Zelenko Z, Aghajanova L, Erikson DW, Barragan F, et al. Perivascular human endometrial mesenchymal stem cells express pathways relevant to self-renewal, lineage specification, and functional phenotype. Biol Reprod. 2012;86(2):58.

30. Kumar A, D’Souza SS, Moskvin OV, Toh H, Wang B, Zhang J, et al. Specification and Diversification of Pericytes and Smooth Muscle Cells from Mesenchymoangioblasts. Cell Reports. 2017;19(9):1902–16.

31. Vento-Tormo R, Efremova M, Botting RA, Turco MY, Vento-Tormo M, Meyer KB, et al. Single-cell reconstruction of the early maternal–fetal interface in humans. Nature. 2018;563(7731):347–53.

32. Shaffer W. Role of Uterine Adhesions in the Cause of Multiple Pregnancy Losses. Clinical Obstetrics and Gynecology. 1986;29(4):912–24.

33. Barragan F, Irwin JC, Balayan S, Erikson DW, Chen JC, Houshdaran S, et al. Human Endometrial Fibroblasts Derived from Mesenchymal Progenitors Inherit Progesterone Resistance and Acquire an Inflammatory Phenotype in the Endometrial Niche in Endometriosis. Biol Reprod. 2016;94(5):118.

34. Kapranov NM, Davydova YO, Galtseva IV, Petinati NA, Drize NI, Kuzmina LA, et al. Effect of Priming of Multipotent Mesenchymal Stromal Cells with Interferon gamma on Their Immunomodulating Properties. Biochemistry (Mosc). 2017;82(10):1158–68.

35. Nemeth K, Leelahavanichkul A, Yuen PS, Mayer B, Parmelee A, Doi K, et al. Bone marrow stromal cells attenuate sepsis via prostaglandin E(2)-dependent reprogramming of host macrophages to increase their interleukin-10 production. Nat Med. 2009;15(1):42–9.

36. Vasandan AB, Jahnavi S, Shashank C, Prasad P, Kumar A, Prasanna SJ. Human Mesenchymal stem cells program macrophage plasticity by altering their metabolic status via a PGE2-dependent mechanism. Scientific reports. 2016;6:38308.

37. Efremova M, Vento-Tormo M, Teichmann SA, Vento-Tormo R. CellPhoneDB: inferring cell–cell communication from combined expression of multi-subunit ligand–receptor complexes. Nature Protocols. 2020;15(4):1484–506.

38. Armulik A, Genové G, Betsholtz C. Pericytes: Developmental, Physiological, and Pathological Perspectives, Problems, and Promises. Developmental Cell. 2011;21(2):193–215.

39. Krueger M, Bechmann I. CNS pericytes: Concepts, misconceptions, and a way out. Glia. 2010;58(1):1–10.

40. The Endometrium. Aplin J, Fazleabas, A., Glasser, S., Giudice, L., editor. London: CRC Press; 2008.

41. Le Blanc K, Davies LC. MSCs-cells with many sides. Cytotherapy. 2018;20(3):273–8.

42. Bradley JE, Ramirez G, Hagood JS. Roles and regulation of Thy-1, a context-dependent modulator of cell phenotype. Biofactors. 2009;35(3):258–65.

43. van den Brink SC, Sage F, Vértesy Á, Spanjaard B, Peterson-Maduro J, Baron CS, et al. Single-cell sequencing reveals dissociation-induced gene expression in tissue subpopulations. Nature methods. 2017;14(10):935–6.

44. Dobin A, Davis CA, Schlesinger F, Drenkow J, Zaleski C, Jha S, et al. STAR: ultrafast universal RNA-seq aligner. Bioinformatics. 2013;29(1):15–21.

45. Butler A, Hoffman P, Smibert P, Papalexi E, Satija R. Integrating single-cell transcriptomic data across different conditions, technologies, and species. Nat Biotechnol. 2018;36(5):411–20.

46. Stuart T, Butler A, Hoffman P, Hafemeister C, Papalexi E, Mauck WM, 3rd, et al. Comprehensive Integration of Single-Cell Data. Cell. 2019;177(7):1888–902.e21.

47. Hafemeister C, Satija R. Normalization and variance stabilization of single-cell RNA-seq data using regularized negative binomial regression. Genome Biol. 2019;20(1):296.

48. Aran D, Looney AP, Liu L, Wu E, Fong V, Hsu A, et al. Reference-based analysis of lung single-cell sequencing reveals a transitional profibrotic macrophage. Nat Immunol. 2019;20(2):163–72.

49. Finak G, McDavid A, Yajima M, Deng J, Gersuk V, Shalek AK, et al. MAST: a flexible statistical framework for assessing transcriptional changes and characterizing heterogeneity in single-cell RNA sequencing data. Genome Biol. 2015;16:278.

50. La Manno G, Soldatov R, Zeisel A, Braun E, Hochgerner H, Petukhov V, et al. RNA velocity of single cells. Nature. 2018;560(7719):494–8.

